# Hamsters with long Covid exhibits a neurodegenerative signature in the brainstem

**DOI:** 10.1101/2024.12.16.628627

**Authors:** Anthony Coleon, Florence Larrous, Lauriane Kergoat, Magali Tichit, David Hardy, Thomas Obadia, Etienne Kornobis, Hervé Bourhy, Guilherme Dias de Melo

**Author notes:** Corresponding author: Guilherme Dias de Melo, Lyssavirus Epidemiology and Neuropathology Unit Institut Pasteur, 25-28, rue du Dr. Roux, 75724 Paris Cedex 15, France.

## Abstract

After infection with SARS-CoV-2, patients may present with one or more symptoms that appear or persist over time, including fatigue, respiratory, cardiovascular and neurological disorders. Neurological symptoms include anxiety, depression and impaired short-term memory. However, the exact underlying mechanisms of long Covid are not yet decrypted. Using the golden hamster as a model, we provide further evidence that SARS-CoV-2 is neuroinvasive and can persist in the central nervous system, as we found viral RNA and replicative virus in the brainstem after 80 days of infection. Infected hamsters presented a neurodegenerative signature in the brainstem, with overexpression of innate immunity genes, impacted dopaminergic and glutamatergic synapses, altered energy metabolism. Finally, the infected hamsters manifested persistent signs of depression and impaired short-term memory, as well as late-onset signs of anxiety, as a valuable model to study long Covid. Conclusively, we provide evidence that virus-related and neurodegenerative and immunometabolic mechanisms coexist in the brainstem of infected hamsters and contribute to the manifestation of neuropsychiatric and cognitive symptoms.

**Highlights:**

- SARS-CoV-2 infects and persists in the brainstem of intranasally-inoculated hamsters
- Persistent neuropsychiatric and cognitive consequences are observed in SARS-CoV-2-infected hamsters
- The brainstem present distinct transcriptome profiles in acute and in long Covid
- The dopaminergic and glutamatergic systems are affected in long Covid
- The SARS-CoV-2 infection affects the expression of genes related to neurodegenerative processes in acute and in long Covid

## Introduction

COVID-19 remains a contemporary global public health issue, with more than 776 million cases reported to date, including more than 7 million deaths ^1^. After the acute phase, not all COVID-19 survivors regain normal health, even several months after the first clinical signs. It’s estimated that 6.5-12% of patients may present persistent symptoms, such as fatigue, anosmia, respiratory problems, cognitive and psychosocial distress, as part of a new entity commonly called long Covid ^2–4^. Long Covid, or post-acute sequelae of COVID-19 (PASC), is a plethora of persisting symptoms experienced by survivors after acute COVID-19, regardless the severity of initial symptoms, even though some risk factors to develop long Covid have been identified, which includes sex, age, body weight and preexisting respiratory affections^5–7^.

SARS-CoV-2, the causative agent of COVID-19, is a neuro-invasive virus and the olfactory bulbs is one of the most likely routes by which it enters the brain, as the virus and inflammation are frequently observed in olfactory mucosa in humans and in animal models^8,9^. SARS-CoV-2 has already been detected in the brain of patients who died from COVID-19, mainly in the brainstem and cranial nerves ^10–12^. In other cases, changes in myelinization and synapse organization were detected in the brainstem and the cerebellum^13^, including signaling defects in specific neuronal populations, including glutamatergic^14^ and dopaminergic ones^15^. In a recent study involving 231 French patients suffering with long Covid, 91.8% of them experience neurological and neurocognitive symptoms one year after their first long Covid consultation ^4^. Viral persistence and a pro-inflammatory status may lead to an altered energetic and neurotransmitter metabolism ^16,17^, all possible factors that could induce the manifestation of neuropsychiatric and cognitive symptoms ^18–21^. However, the exact neuroanatomic basis of these signs is not yet defined.

Here we show that, after the intranasal infection by SARS-CoV-2, infected hamsters exhibits a neurodegenerative signature in the brainstem, which is associated with an active infection and an intense inflammatory response, followed by cellular exhaustion in the late phase, characterized mostly by impacted neurotransmission and altered energy metabolism. All these alterations in the brain which share some hallmarks with neurodegenerative diseases may be the underlying causes of prolonged and persistent neurobehavioral changes, including signs of anxiety, depression, and memory loss. Without social or somatization influence, we describe the hamster model for long Covid and provide conclusive evidence that long Covid is a factual biological issue that follows the acute infection. This work uncovers the long-standing effects of SARS-CoV-2 infection on brain metabolism and behavior.

## Results

### Hamsters infected with SARS-CoV-2 exhibit different clinical profiles in acute and in late phases, with sex- and variant-dependent effects

We performed an 80-days follow-up study (Fig. 1A) to characterize long-term effects of the infection in golden hamsters intranasally-infected with the ancestral SARS-CoV-2 Wuhan and the Delta and Omicron/BA.1 variants. We first investigated the differences in the clinical picture induced by the different viruses in a time- and sex-dependent manner (Fig. 1B-G). Wuhan and Delta induced an important body weight loss and high clinical scores in male hamsters in the acute phase (Fig.1B, C). Female hamsters also exhibited important body weight loss but milder clinical scores compared to males (Fig.1E, F). Animals infected with Omicron/BA.1, however, presented no or very limited weight loss and lower clinical scores compared to animals infected with the other SARS-CoV-2 variants (Fig.1C, F). In summary, when analyzing males and females combined, the severity of the acute disease was Wuhan > Delta > Omicron/BA.1. The peak of the acute phase of the infection was defined as 4 days post-infection (dpi), time-point where the animals presented the highest clinical score (Fig.1C, F). After 10 dpi, all the animals had recovered the weight loss and no more clinical sign was noticeable. From 10 dpi until 80 dpi, the animals continued to gain weight, however, among the males, the infected animals did not reach the weight of the control group by the end of the experiment. At the time of sampling (80 dpi), all animals were considered clinically healthy, nevertheless, the lung weight-to-body weight ratio (LW/BW) was higher in some animals, particularly in male and female hamsters infected with Wuhan, and in male hamsters infected with Omicron/BA.1 (Fig. 1D, G), which indicates that the lungs are heavier, probably due to inflammation, oedema and congestion.

**Fig. 1.**
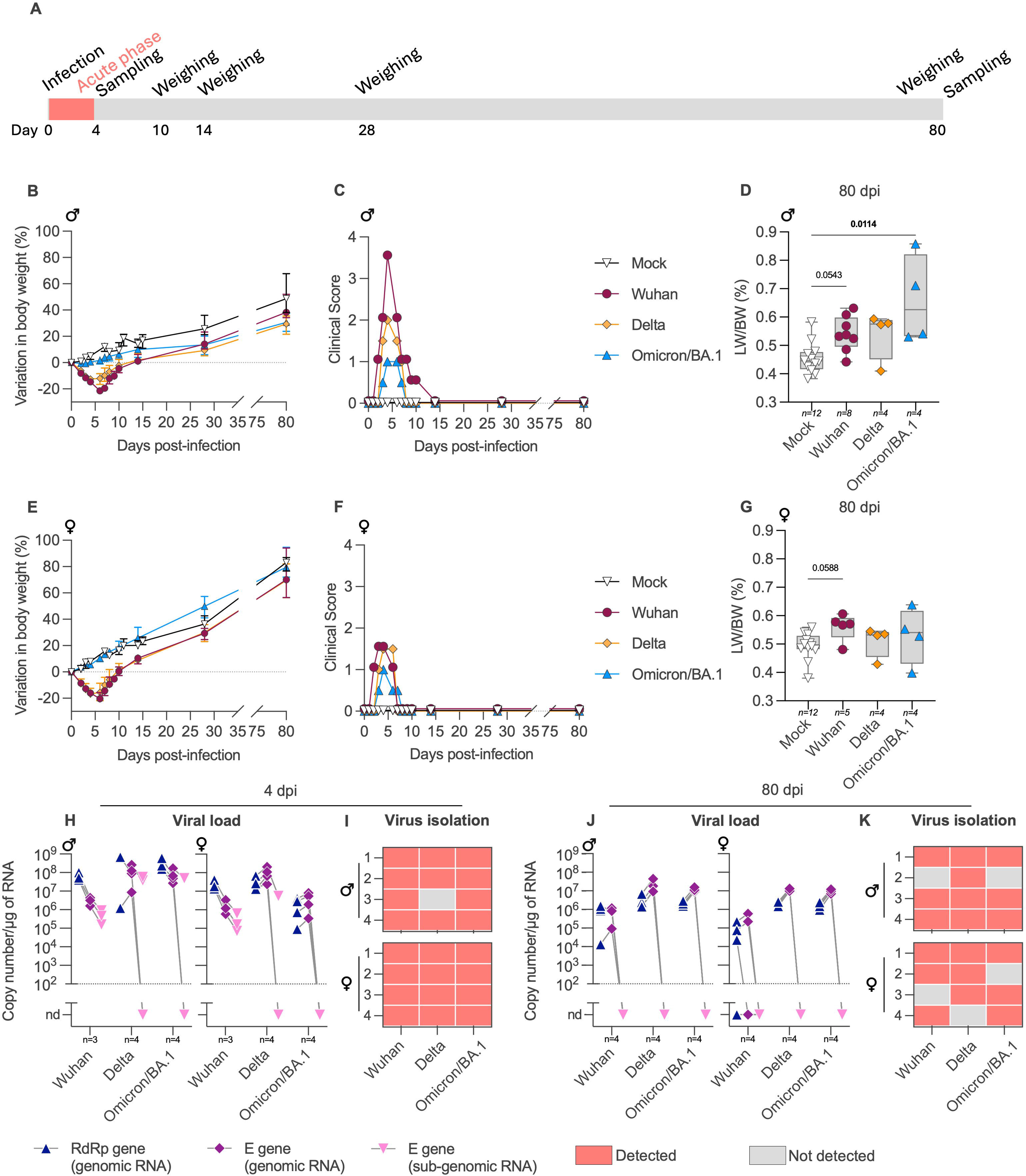
Long-term clinical profile and brainstem viral load in hamsters intranasally-infected with SARS-CoV-2 Wuhan or the variants Delta and Omicron/BA.1. (A) General experimental outline. (B-G) Clinical status in male (♂) and female hamsters (♀LJ). Body weight variation (B,E) and clinical score (C,F) over 80 days post-infection (dpi). The clinical score is based on a cumulative 0-4 scale: ruffled fur, slow movements, apathy, and absence of exploration activity. (D,G) Lung weight-to-body weight (LW/BW) ratio measured at 80 dpi. Kruskal-Wallis test (the adjusted *p* value is shown if *p* < 0.1 and indicated in bold if *p* < 0.05). Horizontal lines indicate the median and the interquartile range. (H-K) SARS-CoV-2 detection in the brainstem at 4 dpi (H-I) and 80 dpi (J-K) in male and female hamsters. (H,J) Genomic and sub-genomic viral RNA were assessed based on the RdRp and E gene sequence. Gray lines connect symbols from the same animals. (I,K) Isolation of infectious virus from the brainstem. Heatmaps showing positive isolation (red squares) or negative isolation (gray squares) of infectious virus. Each square corresponds to one different hamster. The number of animals in each experiment is marked in the graphs. Related to Supplementary Fig.1.

### SARS-CoV-2 infects and persist in the brainstem of intranasally-infected hamsters

We then investigated the presence and persistence of the different SARS-CoV-2 variants in different nervous tissues in relation with the clinical profiles described previously. To this aim, we initially performed a follow-up study to quantify the viral load in the brainstem of intranasally-infected male and female hamsters using Wuhan as model. Remarkably, genomic and sub-genomic viral RNA were detected in the brainstem as early as four hours after the intranasal inoculation, with stable and elevated levels at 1-, 2- and 4-dpi, regardless of the sex. Genomic RNA was still detected at 14-, 30- and 80-dpi, but in contrast, sub-genomic RNA was below the limit of detection (Supplementary Fig. 1A, C).

Next, we measured the viral RNA loads in brainstem of male and female animals infected with Wuhan and the variants Delta and Omicron/BA.1 in two time-points: during the acute infection (4 dpi) and in a late time-point (80 dpi). During the acute phase, SARS-CoV-2 genomic RNA was detected in all samples, regardless of the sex, the variant, or the clinical presentation (Fig. 1H). Sub-genomic RNA was found in all Wuhan-infected samples, and in an inconsistent manner for the Delta- and Omicron/BA.1-infected samples (Fig. 1H). The viral titers in the brainstem were below the quantification limit (∼10^2^ TCID50/mL), nevertheless, we could isolate infectious virus from 96.8% (31/32) of the brainstems from infected hamsters at 4 dpi (Fig. 1I)

Remarkably, we still detected genomic RNA in the brainstem of the animals at 80 dpi (Fig. 1J). Although sub-genomic RNA was below the limit of detection, indicating low viral load, we were able to isolate and amplify infectious virus in 84.4% (27/32) of the brainstems of infected hamsters at 80 dpi (Fig. 1K). In parallel, we also quantified the viral RNA loads in the airways (nasal turbinates and lungs) and in other parts of the brain than the brainstem (olfactory bulbs, cerebral cortex, cerebellum) of the animals at 80 dpi. Genomic SARS-CoV-2 RNA was detected in the airways of all animals. In the other brain areas, however, the detection was variable (Supplementary Fig. 2), and no sub-genomic viral RNA was detected.

### SARS-CoV-2 induces a particular phenotypic glial pattern in the brainstem of infected hamsters

Having demonstrated the persistence of SARS-CoV-2 RNA and infectious virus in different brain areas of infected hamsters, we next analyzed the histopathological changes at 4- and 80-dpi, focusing on the olfactory bulbs (considered the entry point of the virus into the brain) and on the brainstem (Fig. 2A-C). In general, the histopathological analysis of the brain revealed no relevant microscopic alterations neither during the acute nor during the late phase of the infection (Supplementary Fig. 3). In the airways, however, significant lesions were detected at 4 dpi, mostly represented by destruction of the olfactory mucosa, and congestion and inflammation in the lungs (Supplementary Fig. 4).

**Fig. 2.**
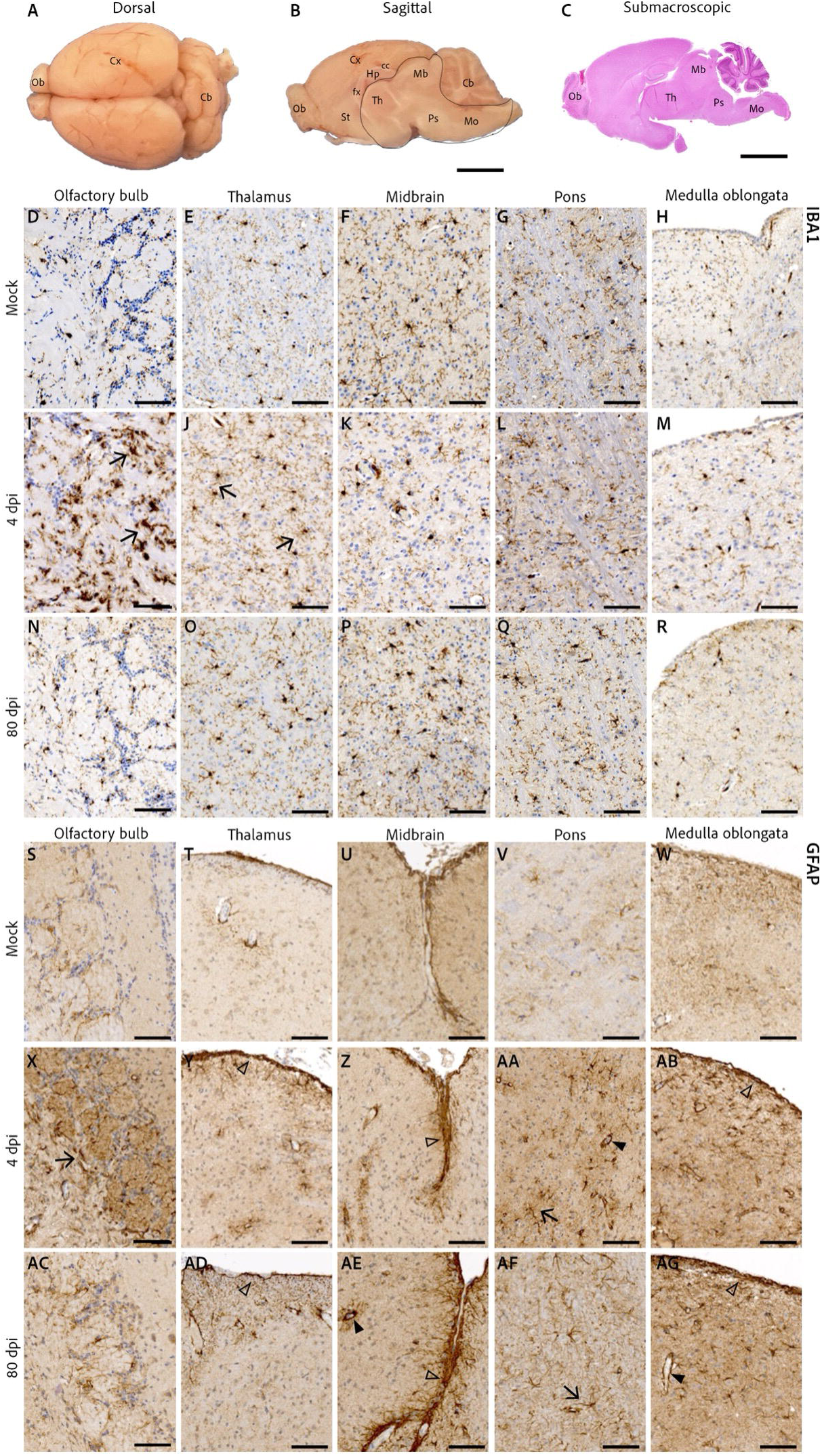
Histopathological analysis in the brain of hamsters intranasally-infected with SARS-CoV-2 Wuhan. (A) Dorsal view of a formalin-fixed hamster brain. Ob: olfactory bulbs, Cx: cortex, Cb: cerebellum. (B) Midsagittal section of a formalin-fixed hamster brain. Hp: hippocampus, cc: corpus calosum, fx: fornix, St: striatum, Th: thalamus, Mb: midbrain, Ps: pons, Mo: medulla oblongata. The brainstem (contoured by a black line) can be separated from the brain after removal of the cerebellum and detachment from the telencephalon at the level of the corpus callosum and the fornix. (C) Submacroscopic view of a sagittal section (HE staining), with indications of the regions from where the images D-R were obtained. (D-H) Microglial detection (IBA1+) in the brain of a mock-infected hamster: olfactory bulb, between the olfactory nerve layer and the glomerular layer (D), thalamus (E), midbrain (F), pons (G) and medulla oblongata (subependymal zone; H). (I-M) Microglial detection in the brain of a hamster at the acute phase of infection (4 days post-infection, dpi), where an intense microglial reactivity (hyperplasia with thicker cell processes) can be observed in the olfactory bulbs (I, arrow) and discrete microgliosis can be randomly found in the thalamus (J, arrow). (N-R) At 80 dpi, microglial cells were in general not reactive. (S-W) Astrocytes (GFAP+) detection in the brain of a mock-infected hamster: olfactory bulb, between the olfactory nerve layer and the glomerular layer (S), thalamus (subependymal zone, T), midbrain at the level of the cruciform sulcus between the superior and inferior colliculus (U), pons (V) and medulla oblongata (subependymal zone; W). (X-AG) In the olfactory bulbs, astrogliosis is evident only at 4 dpi (X, arrow). In the brainstem, however, GFAP-immunoreactive astrocytes are increased in both acute COVID-19 (4dpi, Y-AB) and long Covid (80 dpi, AD-AG), where the *glia limitans superficialis* (subpial astrocyte foot processes; open arrowhead), the *glia limitans perivascularis* (perivascular astrocyte foot processes; filled arrowhead) and parenchymal astrocytes (arrow) are more evident. D-R: Iba-1 IHC to detect microglial cells. S-AG: GFAP IHC to detect astrocytes. Representative images: mock-infected (n=4 males + 4 females), acute COVID-19 (n=4 males), long Covid (n=4 males + 4 females). Scale bars: A-C = 5 mm, D-AG = 100 µm.

To better assess the status of glial cells in the olfactory bulbs and in the brainstem, we used IBA1 (ionized calcium-binding adapter molecule 1) to identify microglial cells (Fig. 2D-R), and GFAP (glial fibrillary acidic protein) to identify astrocytes by immunohistochemistry (Fig. 2S-AG). In the infected group at 4 dpi, we observed foci of reactive microglial cells in the olfactory bulbs, mainly localized between the olfactory nerve layer and the glomerular layer (Fig. 2I), and mild microgliosis in the thalamus (Fig.2J) compared to mock-infected group (Fig. 2D-H.) No distinguishable microglial changes were observed elsewhere at 4dpi (Fig. 2K-M) nor at 80 dpi (Fig. 2N-R). The astrocytic involvement during SARS-CoV-2 presented a slightly different pattern, with higher detection of GFAP (astrogliosis) frequently noticed in the olfactory bulbs, subpial, perivascular and subependymal regions of infected animals at 4 dpi (Fig. 2X-AB) compared to mock-infected hamsters (Fig. 2S-W). Strikingly, subpial, perivascular and subependymal astrogliosis were still evident at 80 dpi (Fig. 2AC-AG).

Finally, to assess neuroaxonal damage, we quantified the levels of neurofilament light chain (NfL) in the serum of male and female hamsters intranasally-infected with Wuhan at 80dpi. No difference was observed, excepted in one male hamster showing an increased level of NfL in the serum when compared with mock-infected control (Supplementary Fig. 1B).

### The brainstem present distinct transcriptome profiles in acute and late infection

Next, willing to unveil if the metabolism of the brainstem was affected during SARS-CoV-2 infection and persistence, we performed a comparative agnostic transcriptomic approach using bulk RNA-seq in the brainstem of hamsters in two distinct comparisons: acute infection (brainstems at 4 dpi in comparison with age-related mock-infected animals), and late infection (brainstems at 80 dpi in comparison with age-related mock-infected animals).

In the brainstem at 4 dpi, we observed 3863 and 4014 differentially expressed genes (DEGs; increased or decreased, respectively), from which 391 and 115 DEGs (increased or decreased, respectively) had a fold change higher than 2. The DEGs were classified according to the GO (gene ontology) terms based on their biological processes, cellular components and molecular functions, and to the KEGG (Kyoto Encyclopedia of Genes and Genomes) pathways. Regarding GO terms, dysregulated biological processes (366 GO terms) were mainly related to inflammation, pyroptosis and cytolysis, type I interferon and viral processes; cellular component (117 GO terms) were related to neurites, synapses and mitochondria, whereas molecular function (55 GO terms) highlighted energy production and channel activity (Fig. 3). Sixty-nine KEGG pathways were significantly regulated, including pathways related to neurodegeneration, neurotransmission, energy, cellular metabolism, autophagy, intracellular signaling, and viral cycle (Fig. 4A).

**Fig. 3.**
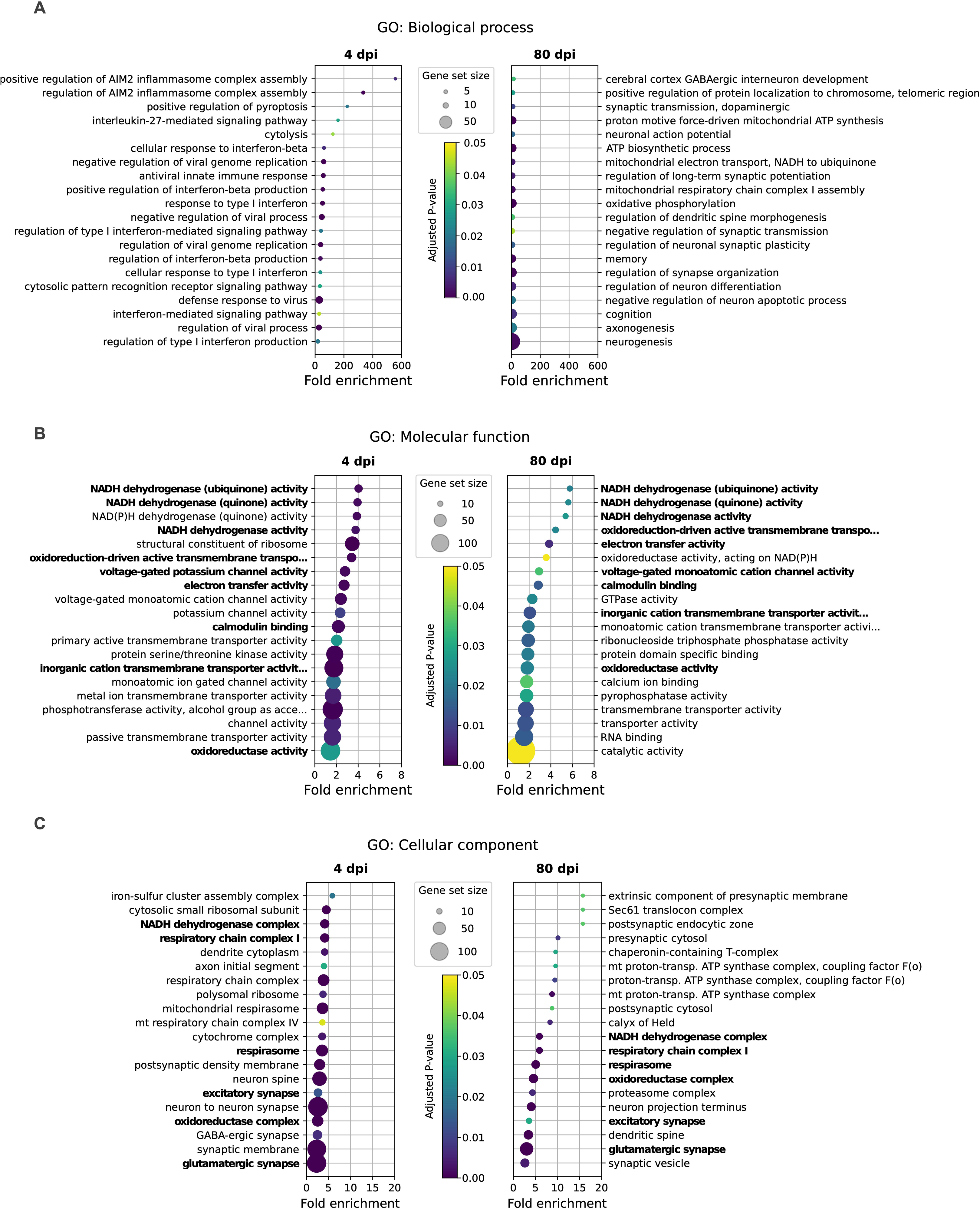
Intranasal SARS-CoV-2 infection alters the brainstem transcriptome profile in hamsters. GO enrichment analysis considering biological process (A), molecular function (B) and cellular compartment (C). Selected GO terms are based on the up- and down-regulated genes between SARS-CoV-2 Wuhan-infected and mock-infected samples at 4 dpi (left panels) and 80 dpi (right panels). Circle sizes are proportional to the gene set size, which shows the total size of the gene set associated with the GO terms. Circle color is proportional to the corrected *p* values. Common pathways observed at 4 and 80 dpi are shown in bold.

**Fig. 4.**
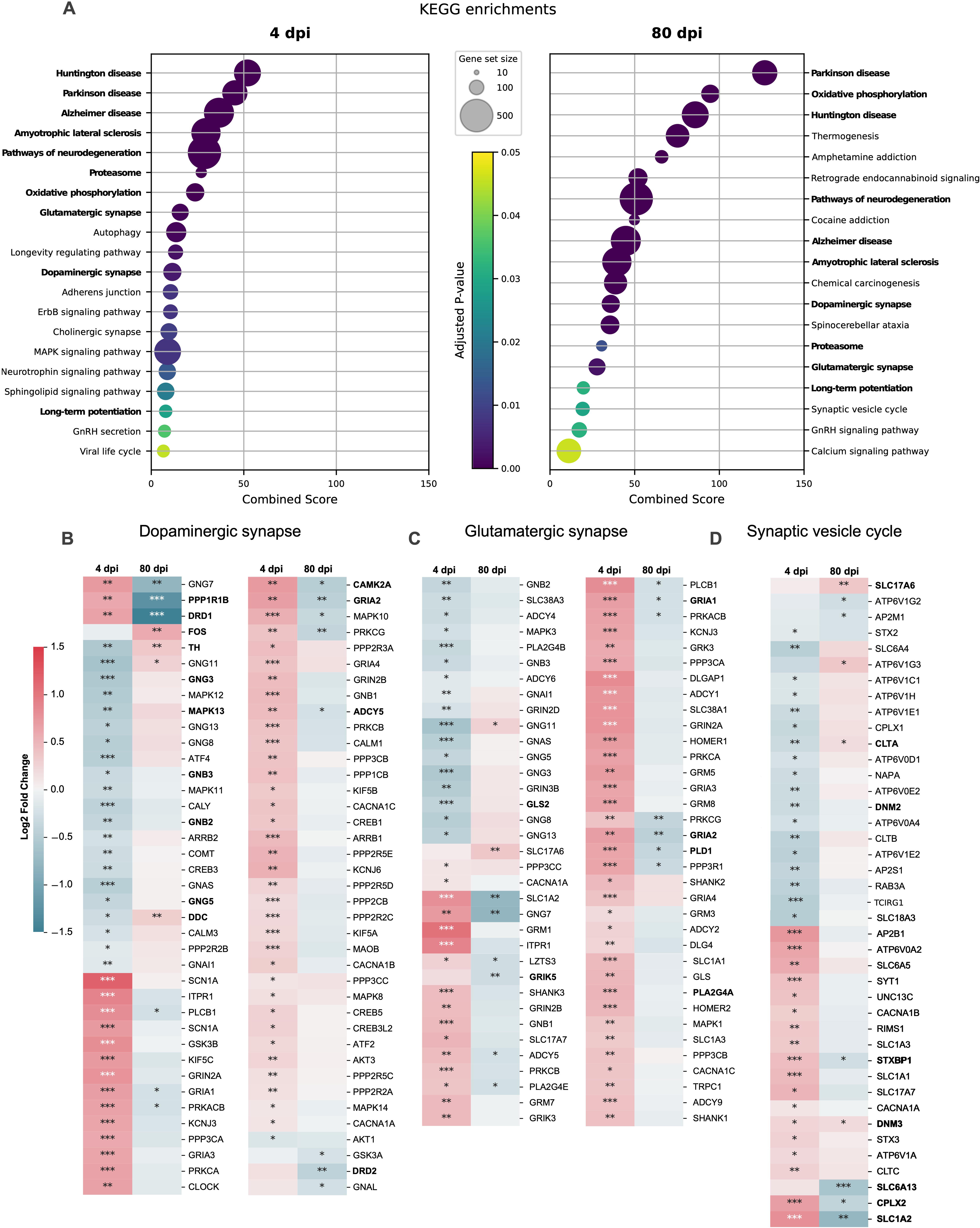
Intranasal SARS-CoV-2 infection alters the brainstem transcriptome profile in hamsters. (A) KEGG pathways enrichment based on the differentially regulated genes between SARS-CoV-2 Wuhan-infected and mock-infected samples at the acute phase (4 days post-infection, dpi; left panel) and at the late phase (80 dpi, right panel). Circle sizes are proportional to the gene set size. Circle color is proportional to the corrected P values. Common pathways observed at 4 and 80 dpi are shown in bold. (B-D) Heatmap showing the differentially expressed genes at 4 and 80 dpi according to the selected KEGG pathway “Dopaminergic synapse” (B), “Glutamatergic synapse” (C), and “Synaptic vesicle cycle” (D), calculated in comparison with mock-infected hamsters. Benjamini–Hochberg-adjusted test, *, **, *** denote *p* < 0.05, *p* < 0.01 and *p* < 0.001 respectively. Color gradient represents the transcription log2 fold change comparing infected and mock-infected. Gene names in bold are mentioned in the text. Related to Supplementary Fig. 5-6.

At 80 dpi, we observed 410 and 424 DEGs (increased or decreased, respectively), from which 7 and 94 DEGs (increased or decreased, respectively) with a fold change higher than 2 in the brainstem. During this late phase of SARS-CoV-2 infection, the dysregulated biological processes in the brainstem (234 GO terms) mainly involved energy, synapses, axonogenesis and neurogenesis; the cellular component (131 terms) was identified mainly as targeting synaptic zones and mitochondria; and molecular function (28 GO terms) was related to energy and calcium activity (Fig. 3). Interestingly, two biological processes terms related to behavior, memory and cognition, were also identified (Fig. 3A). Further, twenty-three KEGG pathways were significantly regulated, mostly related to neurodegeneration, neurotransmission, calcium signaling and energy (Fig. 4A).

Remarkably, several GO terms and KEGG pathways were found to be regulated in the brainstem at both 4 dpi and 80 dpi, but interestingly, the expression of the related genes presented an opposite profile, i.e., most genes found up-regulated at 4 dpi were down-regulated at 80 dpi, and vice-versa (Fig. 4B-D, 5A).

**Fig. 5.**
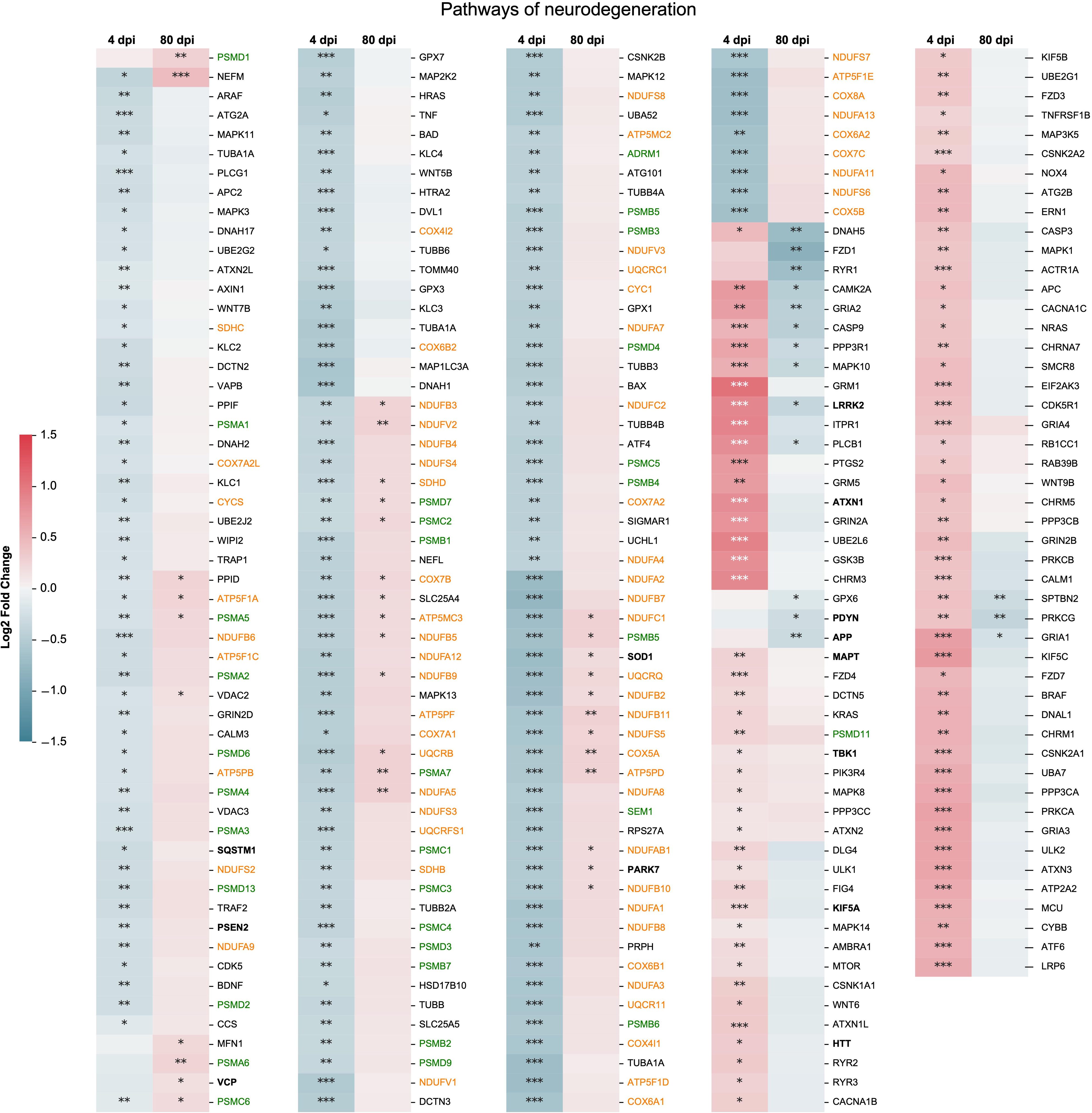
Intranasal SARS-CoV-2 infection affects the expression of genes related to neurodegenerative processes in the brainstem. Heatmap showing the differentially expressed genes at 4- and 80-days post-infection (dpi) according to the selected KEGG pathway “Pathways of neurodegeneration” calculated comparing SARS-CoV-2 Wuhan-infected to mock-infected hamsters. Benjamini–Hochberg-adjusted test, *, **, *** denote *p* < 0.05, *p* < 0.01 and *p* < 0.001 respectively. Commons genes appearing in the ‘Oxidative phosphorylation pathway’ are shown in orange, whereas common genes appearing in the ‘Proteasome pathway’ are shown in green. Color gradient represents the transcription log2 fold change comparing infected and mock-infected. Gene names in bold are mentioned in the text. Related to Supplementary Fig. 6-7.

### The synapse system seems to play a central role in the long-term consequences of Covid-19

Among the common dysregulated pathways observed in the brainstem at 4 dpi and 80 dpi, the dopaminergic and glutamatergic synapses pathways can be highlighted (Fig. 4A). Regarding the dopaminergic pathway, on the one hand, there is a significant reduction in the gene expression related to the enzymes involved in dopamine biosynthesis TH (tyrosine hydroxylase) and DDC (dopa decarboxylase) at 4 dpi, while the genes of dopamine receptors (DRD1) and intracellular mediators (PPP1R1B, also known as DARP-32; CAMK2A) are overexpressed (Fig. 4B). At 80 dpi, on the other hand, there is a decrease of the mRNA expression of dopamine receptors (DRD1, DRD2), AMPA receptors (GRIA1, GRIA2), and intracellular mediators (PPP1R1B, CAMK2A), with an increase of few genes, including FOS, TH and DDC (Fig. 4B).

The glutamatergic synapse pathway followed the same profile as the dopaminergic one, with an upregulation of mRNAs related to glutamate receptors from different categories (AMPA, NMDAR, metabotropic, ionotropic) and down-regulation of key enzymes (glutaminase: GLS2) and different intracellular mediators (GNB2, GNB3, GNG3, GNG5, ADCY5, MAPK3) at 4 dpi. A more restricted number of DEGs was observed at 80 dpi, including glutamate receptors (GRIK5, GRIA2), and intracellular mediators including phospholipases (PLA2G4E, PLD1; Fig.4C).

Further, the synaptic vesicle cycle pathway was also dysregulated, which includes the mRNA expression of key members of the SNARE-complex (Munc18: STXBP1; and complexin: CPLX2), of the clathrin-mediated endocytosis (clathrin: CLTA, CLTB, CLTC; dynamin: DNM2, DNM3) and of neurotransmitter transporters (glutamate: SLC1A2, also known as *EAAT2; SLC17A6, also known as VGLUT2*; GABA: SLC6A13, also known as GAT2; Fig. 4D). Further, the mRNA expression of other neurotransmitter receptors was also impacted in the brainstem of infected hamsters, including downregulation of CHRM4 and HTR2C at 80 dpi (Supplementary Fig 6).

### SARS-CoV-2 infection affects the expression of genes related to neurodegenerative processes in the brainstem

Neurodegeneration was also a term frequently observed in the RNA-seq analyses in the brainstem of hamsters infected with SARS-CoV-2. All KEGG pathways from the database “Neurodegenerative disease” were dysregulated in both at 4- and 80-dpi, namely: Alzheimer disease, Parkinson disease, Amyotrophic lateral sclerosis, Huntington disease, Spinocerebellar ataxia, Prion disease, Pathways of neurodegeneration - multiple diseases (Fig. 4A). The intranasal infection by SARS-CoV-2 dysregulated the expression of a multitude of genes in the brainstem that can contribute, trigger or exacerbate neurodegenerative processes (Fig. 5), including the following seven main processes: *i*: altered energy homeostasis (altered mitochondrial electron transport, complexes I to V) which includes some genes common to the oxidative phosphorylation pathway (shown in orange in Fig. 5). *ii*: synaptic and neuronal network defects (dysregulated synaptic vesicle cycle; abnormal dopaminergic and glutamatergic neurotransmission; Fig. 4B-D). *iii*: aberrant proteostasis (altered proteasome activity, shown in green in Fig. 5). *iv*: genomic instability (dysfunctional base excision repair, nucleotide repair, spliceosome), and *v*: inflammation (more evident in the acute phase; Fig. 3A, Supplementary Fig. 5). Our results further include *vi*: neuronal cell death (loss of trophic support such as BDNF, ErbB signaling) and *vii*: stem cell exhaustion (decreased neurogenesis, axonogenesis).

Additionally, from a list of the highest risk genes to cause neurodegenerative diseases in humans ^22^, we detected ten of them differentially expressed in the brainstem of hamsters at 4 dpi*, ATXN1, HTT, KIF5A, LRRK2* also known as PARK8, MAPT, PARK7 also known as DJ-1, PSEN2, SQSTM1, SOD1, TBK1) and six of them at 80 dpi (APP, LRRK2, PARK7, PDYN, SOD1, VCP).

Lastly, we evaluated the gene expression of selected targets by RT-qPCR in the brainstem of male and female hamsters infected with Wuhan, Delta and Omicron/BA.1 at 4 and 80 dpi to corroborate the involvement of the abovementioned signaling pathways identified by RNA-seq. The genes MX2, ISG20, IFNL and CXCL10 were upregulated in the brainstem of both male and female hamsters at 4 dpi, in agreement with the pathways related to inflammation and antiviral innate immune response in the acute phase (Supplementary Figure 5). Regarding dopaminergic ang glutamatergic synapses impairment, the genes DRD2, FOS, CAMK2A, PLA2G4E, CHRM4, GRID1 and SOX2 were found upregulated in the brainstem of males and females at 4 dpi, whereas DRD1 and ADCY5 were upregulated only in females. Differences in the gene expression were less intense at 80 dpi, nevertheless, we observed a downregulation of HTR2C in the brainstem of both males and females; downregulation of DRD1, PLA2G4 and GRID1 in females; and downregulation of DRD2 and CAMK2A in males (Supplementary Fig. 6). Concerning neurogedeneration, oxidative phosphorylation, and proteasome pathways at 4 dpi, PSMB5 expression was reduced in male brainstems; CHRM1 and ATP5ME were upregulated in female brainstems. In contrast, at 80 dpi, PSMB5 was upregulated in the brainstem of both sexes, whereas in males, COX5A and ATP5ME were upregulated and CHRM1 was downregulated (Supplementary Fig. 7).

### SARS-CoV-2-infected hamsters manifest long-term neuropsychiatric symptoms and cognitive deficit

Finally, to check if the altered brainstem metabolism due to SARS-CoV-2 infection would be enough to cause clinical manifestations related to long Covid, three well described and recognized behavioral tests were performed to assess anxiety, depression, and recognition memory in the animals. The hamsters performed four sequential behavioral tests (one test/day) on three sessions: the first session just after the acute phase (between 14-17 dpi, hereinafter called 15 dpi session), the second between 28-31 dpi (hereinafter called 30 dpi session), and the third session between 76-79 dpi (hereinafter called 80 dpi session) (Fig. 6A). To properly quantify the effect of covariates (sex, time post-infection, SARS-CoV-2 variant), we used mixed model regression to account for within-individual correlation arising from repeated measurements.

**Fig. 6.**
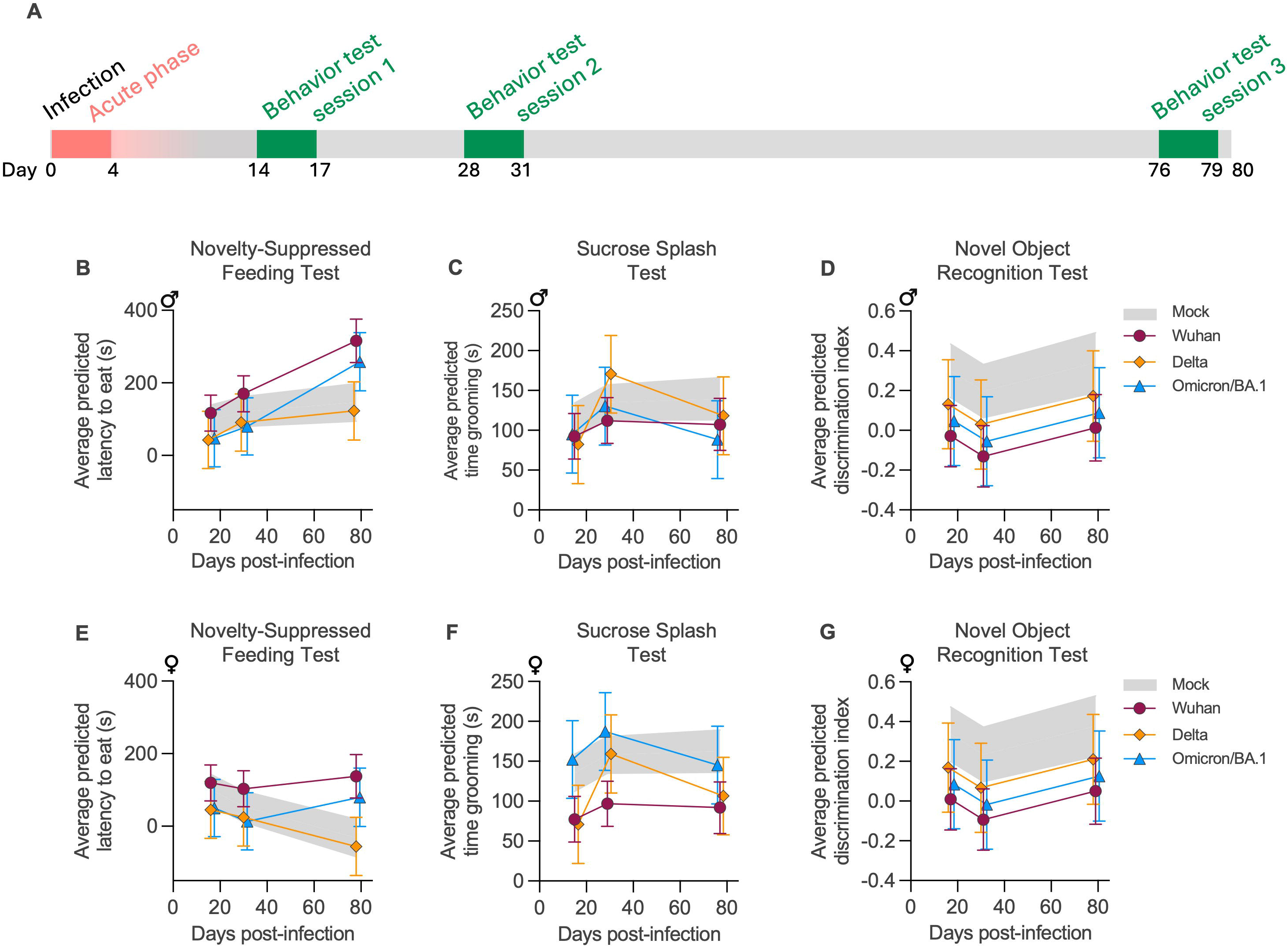
Long-term impact of SARS-CoV-2 infection on neuropsychiatric and cognitive behavior in hamsters. (A) General experimental outline with the three sessions of behavior testing. (B,E) Novelty-suppressed feeding test: Average predicted latency to eat analyzed by mixed model regression in male (B) and female (E) hamsters. Increased latency to eat indicates hyponeophagia and corresponds to an anxious behavior. (C,F) Sucrose splash test: Average predicted time grooming analyzed by mixed model regression in male (C) and female (F) hamsters. Time grooming corresponds to an index of motivational and self-care behavior and decreased grooming time is considered as a depression-like behavior. (D,G) Novel object recognition test: Average predicted discrimination index analyzed by mixed model regression in male (D) and female (G) hamsters. Decreased discrimination index corresponds to less time spent exploring the new object and is indicative of short-term memory impairment. Data were analyzed using a mixed-model regression based on time post-infection, sex, and viral variant. Horizontal lines indicate the mean and the 95% confidence interval. The gray crosshatched zone indicates the mean and the 95% confidence interval of the mock-infected group. Related to Supplementary Fig. 8-11.

First, we evaluated signs of anxiety by using the well-known light-dark box and the novelty suppressed feeding tests. During the novelty suppressed feeding test, infected animals displayed anxiety-like behavior, referred to as hyponeophagia (i.e. inhibition of feeding caused by a new environment: anxiety behavior = increased latency to eat) at the 80 dpi session (Fig.6B,E). This phenomenon was more pronounced in male hamsters infected with SARS-CoV-2 Wuhan, and to a lesser extent with Omicron/BA.1 (Supplementary Fig. 8). Regarding the light-dark box (which is based on rodents’ innate aversion to light), this test was unable to differentiate anxious behavior among the groups (Supplementary Fig. 9), which may have been affected by different factors, including the robustness of the test and the lighting conditions of the BSL-3 isolators room.

Next, we assessed depression by carrying out the sucrose splash-test. This test consists of comparing the time that the animals spend grooming after spraying a 10% sucrose solution on their backs: the time grooming corresponds to an index of motivational and self-care behavior (decreased time grooming indicates depression-like behavior). The test showed a dichotomy between males and females: Wuhan-infected females presented a persistent depressive behavior (15, 30, and 80 dpi sessions), whereas Delta-infected females presented a relapsing depressive behavior (15 and 80 dpi sessions) (Fig. 6C, F). Depression incidence was lower for Omicron/BA.1-infected females (Supplementary Fig. 10).

Lastly, to assess the short-term memory capacity of the infected animals, we used the novel object recognition test, where a low discrimination index indicates impaired recognition memory. Strikingly, both male and female animals infected with Wuhan presented persistent memory impairment at all sessions (15, 30, and 80 dpi; Fig. 6D, G), whereas hamsters infected with the Delta and Omicron/BA-1 variants displayed more variable results (Supplementary Fig. 11).

## Discussion

Long Covid is a complex condition that can affect different organs and that can be manifested as a multitude of symptoms after COVID-19. Since the first reports by patients, there have been different definitions of long Covid, and the most recent one was proposed by the United States National Academies of Sciences, Engineering, and Medicine (NASEM) as “an infection-associated chronic condition that occurs after SARS-CoV-2 infection and is present for at least 3 months as a continuous, relapsing and remitting, or progressive disease state that affects one or more organ systems” ^23^. However, despite a well-accepted definition, and the existence of different hypothesis and evidence ^24–26^, the exact underlying mechanisms of long Covid are not yet decrypted, especially those related to the manifestation of neurologic and neuropsychiatric symptoms. Here we describe clinical, behavioral, virologic and transcriptomic data in the golden hamster, a relevant model to study long Covid. Without social or somatization influence, and no effects related to post-intensive care syndrome (PICS), we provide conclusive evidence that long Covid is a factual biological issue that follows the acute infection.

The hamster model presented herein recapitulates long Covid in humans. In human patients living with long Covid, the incidence of cognitive and neuropsychiatric symptoms may vary according to different studies ^27–29^ : in a meta-analysis regrouping 10,530 patients from 18 studies, 32% of them had brain fog and 28% had memory impairment ^30^. Another study reports that among 9,605 patients living with long Covid, 22% had anxiety and 21% manifested signs of depression ^31^. Remarkably, a recent study with human volunteers that were infected by the original SARS-CoV-2 strain showed that the infection led to a persistent deficit in recognition memory and spatial planning, even if the infected volunteers did not report subjective cognitive deficit ^32^.

Our long-term and sequential behavioral testing approach revealed that infected hamsters, despite sex- and SARS-CoV-2 variant-related variations, also showed persistent signs of depression and a deficit in recognition memory, as well as delayed signs of anxiety after acute infection, which can persist or fluctuate overtime. These cognitive and neuropsychiatric symptoms can have different origins and causes; however, the asset of the hamster model is that the behavioral tests took place in an environment free from social pressure or risk of somatization, i.e., infected and mock-infected animals (both male and female) were tested simultaneously under the exact same conditions. Consequently, the differences observed in clinical profile between the infected and mock-infected animals can only be explained by the SARS-CoV-2 infection itself, and the underlying mechanisms are likely to be associated with virus-related and neuro-immunometabolic changes in their brainstem.

Virus-related mechanisms include SARS-CoV-2 persistence, either infectious (replicative) viral particles or viral components in the brain. The neuroinvasiveness of SARS-CoV-2 ^33^ has already been shown in post-mortem samples of human COVID-19 ^10,12^ and reproduced in animal models whose main focus was the olfactory pathway ^9,34,35^. Herein, we show that SARS-CoV-2 quickly reaches the brainstem after intranasal inoculation, with an active replication of all tested variants (Wuhan, Delta, Omicron/BA.1) during the acute phase of the infection, which corroborates other studies reporting brainstem infection in human cases of acute COVID-19 ^10,11^. Surprisingly, we report the unprecedented isolation of infectious viral particles from the brainstem of hamsters with long Covid 80 days after infection, regardless of sex and SARS-CoV-2 variant. This contradicts the common assumption that there are no infectious virus in the brain ^36,37^. Also, it emphasizes the need to use later time points than only 30 days when working with small animals to reproduce long Covid. The infectious viral titer in the brainstem of the hamsters was however quite low in both acute and late phases of the infection; we could not define if the virus was restricted to specific areas or in specific cell populations, nevertheless, the acute and persistent low-level SARS-CoV-2 infection in the brainstem could, at least partially, contribute to the occurrence of long Covid.

Concomitantly to virus-related mechanisms, we found significant neuro-immunometabolic changes in the brainstem of the infected hamsters, that can be divided into three major categories: inflammation, neurotransmission, and energy. Signs of inflammation in the brainstem of infected hamsters were better observed during the acute phase of the infection, where an antiviral response characterized by the expression of IFN-related genes (ISG15, MX2, IRF7) and a focal and discrete glial activation occurs. Such immune profile recapitulates what was demonstrated in humans ^12,13^ and in other brain regions of hamsters ^9^, however, inflammation may last longer ^13,38,39^.

Contrastingly, neurotransmission seems to be greatly affected during both acute and long Covid, and infected hamsters presented a transcriptome profile highly indicative of dopaminergic and glutamatergic impairment. We show a persistent alteration in dopaminergic signaling with dysregulation in the gene expression of key enzymes related to dopamine synthesis (tyrosine hydroxylase, DOPA decarboxylase) and dopamine receptors (DRD1, DRD2). Most of the dopaminergic neurons in the mammal brain resides in the midbrain (part of the brainstem), are the main source of dopamine ^40^ and play an essential role in behavior and the reward system ^41^. Concordantly, the density of tyrosine hydroxylase-reactive neurons was reduced in in patients who died from acute COVID-19^15^, and tyrosine hydroxylase-reactive neurons and neurites in the midbrain were found positive to SARS-CoV-2 nucleocapsid ^10^. Finally, previous studies showed that dopaminergic neurons are permissive to SARS-CoV-2 *in vitro*, and that the infection lead to an inflammatory response and cellular senescence ^15^, and that the infection could exacerbate Parkinson’s disease by inducing dopaminergic neuron death ^42^. With the hamster model described herein, we establish that the perturbation of the dopaminergic network may persist during a long period after the acute phase of the disease.

Parallelly to dopamine, impacted glutamate signaling may also play a role in mental fatigue, mood disorders and cognition deficits ^43–45^, and it was recently described that some dopaminergic neurons in the midbrain can also co-transmit glutamate ^46,47^. Glutamate is the main excitatory neurotransmitter in the brain ^48^ and it’s involved in learning and memorization ^49^. Besides neurons, astrocyte dysfunction may further impair glutamatergic signaling and be involved in the pathophysiology of long Covid ^14^ ; astrocytes are responsible for most glutamate removal from the synaptic cleft via the receptor SLC1A2 (also known as glutamate transporter 1 - GLT1 or excitatory amino acid transporter 2 -EAAT2) ^44^, found down-regulated in the brainstems of the hamsters with long Covid. Finally, we cannot exclude the involvement of other neurotransmission pathways, as GABAergic and cholinergic impacts have equally been reported during SARS-CoV-2 infection ^50,51^ . In agreement with other studies, the results presented herein also advocate for a broader impact on synapses, either via dysregulation of synaptic members gene expression ^52^, destruction or degeneration of synaptic circuits ^53^, and even physical obstruction to the neurotransmission due to changes in synapse morphology ^52^, to viral-induced neuronal fusion ^54^, neuroaxonal damage ^55^ or even loss of myelin ^56^.

Along with impaired neurotransmission, energy metabolism and mitochondrial dysfunction may also be involved in neuropsychiatric pathologies, including depression ^57^. Intriguingly, zones of hypometabolism in various brain areas, including the brainstem, have been identified by [^18^F]fluorodeoxyglucose (FDG)-PET in patients living with long COVID ^58–61^. Additionally, a magnetic resonance imaging study including 401 patients with positive diagnosis of SARS-CoV-2 infection revealed an important difference in the volume of the brainstem, besides tissue damage in areas functionally connected to the primary olfactory cortex ^62^ . In order to maintain synaptic activity, mitochondrial function and energy supply must be tightly controlled, and deregulation of these processes can result in neurodegenerative damage ^22^. A number of signaling pathways associated with neurodegenerative diseases were found to be affected in the brains of patients with long Covid, including Alzheimer’s ^63^ and parkinsonism ^64^, probably by accumulating α-synuclein aggregates ^65^. The structural, metabolic, and synaptic changes observed in our data are also hallmarks of neurodegenerative diseases, including the expression of genes highly correlated to neurodegeneration in humans ^22^.

The upregulation of innate immunity genes and dowregulation of energy metabolism and proteostasis genes have been appointed as a pan-neurodegenerative gene signature, implying neuronal function loss ^66^. Altogether, the results presented herein argues for a neurodegenerative signature in the brainstem induced by SARS-CoV-2 infection, a characteristic shared by some human neurodegenerative diseases ^67,68^ and which may explain the long-term symptoms observed during long Covid ^38,39,69^. Although we do not advocate that SARS-CoV-2 causes neurodegenerative diseases, our findings highlight that the infection disrupts important metabolic pathways, already in the acute phase, which can trigger neurodegenerative processes and neurobehavioral changes, as may be the case in other viral encephalitis ^16,17,70^.

Health agencies and patients’ associations alert that patients’ access to appropriate care is not always well organized, that health professionals are not well informed and that there may be a tendency towards “stigma” and “psychiatrization” of symptoms, where healthcare professionals quickly categorize long Covid symptoms as somatoform ^71,72^. In this study, we describe hamsters as a robust animal model of long Covid and provide evidence that virus-related and neuro-immunometabolic mechanisms coexist in the brainstem, which exhibit a neurodegenerative signature characterized by elevated innate immune response and impacted neurotransmission and energy metabolism. Conclusively, we demonstrate that golden hamsters develop persistent and late-onset neurological symptoms after SARS-CoV-2 infection as a valuable model to study long Covid.

## Material and methods

### SARS-CoV-2 virus and variants

The SARS-CoV-2 Wuhan BetaCoV/France/IDF00372/2020 isolate (EVAg collection, Ref: 014V-03890) was obtained from the National Centre for Respiratory Viruses (Paris, France). The SARS-CoV-2 Delta/2021/I7.2 200 isolate (GISAID ID: EPI_ISL_2029113) and the SARS-CoV-2 Omicron/B1.1.529 isolate (Omicron/BA.1, GISAID ID: EPI_ISL_6794907) were given by the Virus and Immunity Unit of the Institut Pasteur.

Viral stocks were produced on Vero-E6 cells at an MOI (Multiplicity of Infection) of 10^-4^. After three days of infection, the supernatant was collected, clarified and aliquoted for long-term storage at -80°C. Viral stocks were titrated on Vero-E6 by classical plaque assays using semisolid overlays (Avicel, RC581-NFDR080I) ^73^.

### Animal study design and infection

All animal experiments were performed according to the French legislation and in compliance with the European Communities Council Directives (2010/63/UE, French Law 2013–118, February 6, 2013) and according to the regulations of Pasteur Institute Animal Care Committees. The Animal Experimentation Ethics Committee (CETEA 89) of the Institut Pasteur approved this study (200023; APAFIS#25326-2020050617114340 v2) before experiments were initiated.

Male and female golden hamsters (*Mesocricetus auratus;* RjHan:AURA) of 5-6 weeks of age (average weight 60-80 grams) were purchased from Janvier Laboratories (Le Genest-Saint-Isle, France) and handled under specific pathogen-free conditions. The animals were housed by groups of 4 and manipulated in isolators in a biosafety level-3 facility in the Institut Pasteur’s animal facilities accredited by the French Ministry of Agriculture for performing experiments on live rodents. The animals had *ad libitum* access to water and food. Shredded cardboard and a plastic ball were placed on the bedding of each cage in the spirit of promoting refinement. Before any manipulation, animals underwent an acclimation period of one week.

Following the 3Rs principle in animal experimentation, some samples used in this study are originated from animals that also took part in other studies, where clinical status and respiratory viral load were already published ^35,74^. Animals were anesthetized with an intraperitoneal injection of 200 mg/kg ketamine (Kétamine 1000, Virbac France) and 10 mg/kg xylazine (Rompun, Bayer), and 100 µL of physiological solution containing 6×10^4^ PFU of SARS-CoV-2 was administered intranasally to each animal (50 µL/nostril). Mock-infected animals received the physiological solution only. Infected and mock-infected hamsters were housed in separate isolators and were followed-up daily which the body weight and the clinical score were noted. The clinical score was based on a cumulative 0-4 scale: ruffled fur, slow movements, apathy, absence of exploration activity.

### Tissue sampling

At 4 hours, 1-, 2-, 4-, 14-, 30- and 80-days post-infection, the animals were euthanized with an excess of anesthetics (ketamine and xylazine) and exsanguination ^75^. The brain was extracted from the skull, the two brain hemispheres were separated by a median incision and macroscopically divided in four regions using tweezers: (1) olfactory bulbs, (2) cerebellum, (3) cerebral cortex (containing the cortex, the striatum and the hippocampus) and (4) brainstem (containing the diencephalon, the midbrain, the pons and the medulla oblongata) (Figure 2). The lungs were extracted from the thorax and weighted. The nasal turbinates were extracted by opening of the nasal cavity after incision of the nasal and frontal bones. The samples were immediately frozen at -80°C.

Frozen samples were weighed and transferred to Lysing Matrix M 2 mL tubes (116923050-CF, MP Biomedicals) containing 1 mL of ice-cold DMEM (Dulbecco’s Modified Eagle medium, Gibco) supplemented with 1% penicillin/streptomycin (15140148, Thermo Fisher). Samples were homogenized using the FastPrep-24™ system (MP Biomedicals), and the following scheme: homogenization at 4.0 m/s for 20 sec, incubation at 4°C for 2 min, and further homogenization at 4.0 m/s for 20 sec. The tubes were centrifuged at 10,000 × g during 2LJmin at 4LJ°C, and the supernatants collected and stored at -80°C until further analysis.

### Serum sampling and neurofilament light chain (NfL) dosing

At 80-days post-infection, male (n=4) and female (n=4) hamsters infected with SARS-CoV-2/Wuhan, and age-related male (n=4) and female (n=4) mock-infected hamsters were anesthetized (ketamine and xylazine) and blood was collected by intracardiac punction before euthanasia. The blood was store in 1.5 mL tubes and allowed to coagulate at room temperature during at least 30 minutes. The tubes were centrifuged at 2,000 × g during 10LJmin at 4LJ°C, and the serum was collected and stored at -80°C. Before NfL dosing, the serum samples (45 µL) were treated with 1% (v/v) Triton X-100 (5 µL) at room temperature for 2 h. The serum levels of NfL were measured using the commercially available single molecule array (SIMOA) assay NF-Light v2 Advantage kit (Quanterix, 104073) on an HD-X analyzer following the manufacturer’s instructions.

### RNA extraction

Total RNA from lungs, cerebral cortex, cerebellum and brainstem was extracted using the Direct-zol RNA MiniPrep kit (R2052, Zymo Research). Total RNA from nasal turbinates and olfactory bulbs was extracted using the Direct-zol RNA MicroPrep kit (R2062, Zymo Research). In both cases, 125 µL of tissue homogenate were incubated with 375 µL of Trizol LS (10296028, Invitrogen) and the extraction was performed according to the manufacturer’s instructions.

### SARS-CoV-2 detection in golden hamsters’ tissues

The detection of genomic and sub-genomic SARS-CoV-2 RNA was based on the RdRP and the E genes ^76^. We used the SuperScript III Platinum One-Step qRT-PCR kit (Invitrogen 11732-020) in a final volume of 12.5LJμL per reaction in 384-wells PCR plates using a thermocycler (QuantStudio 6 Flex, Applied Biosystems). Briefly, 2.5LJμL of RNA were added to 10LJμL of a master mix containing 6.25LJμL of 2X Reaction mix, 0.2LJµL of MgSO4 (50LJmM), 0.5LJµL of Superscript III RT/Platinum Taq Mix (2 UI/µL), 0.025 µL of ROX Reference and 3.025LJμL of nuclease-free water containing 400LJnM of primers and 200LJnM of probe. To detect the genomic RNA, we used the IP2_primers and probe (IP2_FW 5’-ATGAGCTTAGTCCTGTTG-3’; IP2_RV 5’-CTCCCTTTGTTGTGTTGT–3’ ; IP2_Probe FAM-AGATGTCTTGTGCTGCCGGTA-TAMRA, and the E_sarbeco primers and probe (E_Sarbeco_F1 5′-ACAGGTACGTTAATAGTTAATAGCGT-3’; E_Sarbeco_R2 5′-ATATTGCAGCAGTACGCACACA-3’; E_Sarbeco_Probe FAM-5′-ACACTAGCCATCCTTACTGCGCTTCG-3’-TAMRA). The detection of sub-genomic SARS-CoV-2 RNA was achieved by replacing the E_Sarbeco_F1 primer by the CoV2sgLead primer (CoV2sgLead-Fw 5′-CGATCTCTTGTAGATCTGTTCTC-3’). A synthetic gene encoding the PCR target sequences was ordered from Thermo Fisher Scientific. A PCR product was amplified using Phusion™ High-Fidelity DNA Polymerase (Thermo Fisher Scientific) and *in vitro* transcribed by means of Ribomax T7 kit (Promega). RNA was quantified using Qubit RNA HS Assay kit (Thermo Fisher scientific), normalized, and used as a standard to quantify RNA absolute copy number. The amplification conditions were as follows: 55LJ°C for 20LJmin, 95LJ°C for 3LJminutes, 50 cycles of 95LJ°C for 17LJs and 58LJ°C for 30LJs; followed by 40LJ°C for 30LJs.

### Isolation and amplification of SARS-CoV-2 from the brainstem

Initially, we used classical TCID50 method on Vero-E6 cells to quantify the infectious virus load in the brainstem ^77^. Briefly, serial dilutions of homogenized organs (1:5 up to 1:640) were made in DMEM (Dulbecco’s Modified Eagle medium, Gibco) supplemented with 1% Penicillin/Streptomycin (15140148, Thermo Fisher) and 1 µg/mL TPCK. We added 100 µL of each dilution to 6 wells of a 96-well plate. Next, 100 µL of a preparation containing 8×10^5^ Vero-E6 per mL of DMEM supplemented with 1% Penicillin/Streptomycin and 1µg of TPCK is added. The plate was placed in the incubator at 37°C, 5% CO_2_ for 3 days.

Viral titers were below the quantification limit (∼10^2^ TCID50/mL), although some wells in the titration plates presented cytopathic effects. In an attempt to amplify the virus, the supernatant from lysed (totally or partially) wells was recovered, diluted (1:10) and 1mL/well was added in a 6-well plate containing 1×10^6^ Vero-TMPRSS2 cells. After one hour, the supernatant was removed and replaced by 2 mL of DMEM supplemented with 1% Penicillin/Streptomycin and incubated during 3 or 7 days, for brainstem samples collected at 4 or 80 dpi respectively. The cells were then lysed with Trizol LS (10296028, Invitrogen), the RNA was extracted with the Direct-zol RNA MicroPrep kit (R2062, Zymo Research) and RT-qPCRs were performed with the IP2 and E primers and probes as described above.

### Transcriptomics analysis in golden hamsters’ brainstems

Two RNA-seq studies were conducted: (1) at 4 dpi (acute COVID-19), the infected group was composed of 4 male and 4 female hamsters infected with Wuhan and collected at 4 dpi, whereas the mock-infected group was composed of 3 male and 3 female mock-infected hamsters with samples processed at the same time; and (2) at 80 dpi (long Covid), the infected group was composed of 4 male and 4 female hamsters infected with Wuhan and collected at 80 dpi whereas the mock-infected group was composed of 4 male and 4 female mock-infected hamsters processed at the same time.

RNA preparation was used to build libraries using a Illumina Stranded mRNA library Preparation Kit (Illumina, USA) following the manufacturer’s protocol. Quality control was performed on an Agilent BioAnalyzer. Illumina NovaSeq X sequencer was used to produce paired-end 150b reads from the sequence libraries.

The RNA-seq analysis was performed with Sequana 0.15.3 ^78^. We used the RNA-seq pipeline 0.17.2 (https://github.com/sequana/sequana_rnaseq) built on top of Snakemake 7.32.4 ^79^. Briefly, reads were trimmed from adapters using Fastp 0.22.0^80^ then mapped to the golden hamster MesAur1.0.100 genome assembly from Ensembl using STAR 2.7.10b ^81^. FeatureCounts 2.0.1 ^82^ was used to produce the count matrix, assigning reads to features using the corresponding annotation from Ensembl with strand-specificity information. Quality control statistics were summarized using MultiQC 1.11 ^83^. Statistical analysis on the count matrix was performed to identify differentially regulated genes. Clustering of transcriptomic profiles were assessed using a Principal Component Analysis (PCA). Differential expression testing was conducted using DESeq2 library 1.34.0 ^84^ scripts indicating the significance (Benjamini-Hochberg adjusted p-values, false discovery rate FDR < 0.05) and the effect size (fold-change) for each comparison.

Finally, enrichment analysis was performed using modules from Sequana. The GO (Gene Ontology Consortium 2021) enrichment module uses PantherDB ^85^ and QuickGO ^86^ services. The KEGG pathways enrichment uses GSEApy 1.0.4 ^87^, EnrichR ^88^ and KEGG database ^89^. Programmatic accesses to online web services were performed via BioServices 1.11.2 ^90^.

### Quantitative PCR from golden hamsters’ brainstems during long Covid

To validate the RNA-seq results, 25 genes from the main representative KEGG pathways were selected for quantifying host transcripts in brainstem samples from male and female hamsters infected by Wuhan or by the variants Delta and Omicron/BA.1 collected at 80 days post-infection. Briefly, RNA was reverse transcribed to first strand cDNA using the SuperScript™ IV VILO™ Master Mix (11766050, Invitrogen). qPCR was performed in a final volume of 10LJμL per reaction in 384-well PCR plates using a thermocycler (QuantStudio 6 Flex, Applied Biosystems) and its related software (QuantStudio Real-time PCR System, Design and Analysis Software v. 2.7.0, Applied Biosystems). In each well, 2.5LJμL of cDNA (12.5LJng) were added to 5LJμL of Power SYBR Green PCR Master Mix (4367659, Applied Biosystems) and 2.5LJμL of nuclease-free water containing 1 µM of golden hamster’s primer pairs (Supplementary Table 1). The amplification conditions were as follows: 50LJ°C for 2LJmin, 95°C for 10 min, and 45 cycles of 95LJ°C for 15LJs and 60LJ°C for 1LJmin, followed by a melt curve from 60LJ°C to 95LJ°C. The ACTB and the HPRT genes were used as reference. Variations in gene expression were calculated as the n-fold change in expression in the tissues from the infected hamsters compared with the tissues of the mock-infected group using the 2^-ΔΔCt^ method ^91^.

### Histopathology and immunohistochemistry

Samples destinated to histopathology were collected and processed as follows: the brains were extracted from the skull and fixed in one piece. To obtain intact nasal turbinates, the head of the animals were fixed in one piece after removal of the brain, the mandible and the skin. After fixation, the fragments were incubated with Osteomoll solution (101736, Merck) for 1 day followed by EDTA 0.5M for 4 days to allow bone decalcification; afterwards, the nasal fragments were sectioned either in sagittal, in transversal (Figure 4). All samples were fixed in 10% neutral-buffered formalin for 24-48 hours and embedded in paraffin. Four-µm-thick sections were cut and stained with hematoxylin and eosin staining. Sections were cut to 4 µm thickness and processed for routine histology using hematoxylin and eosin staining and choromogenic ImmunoHistoChemistry (IHC) with hematoxylin counterstaining. IHC analysis was performed using a Rabbit-Iba-1 (019-19741, Wako chemical) and Chicken-GFAP (ab4674, Abcam) antibodies. All slides were scanned using the AxioScan Z1 (Zeiss) system, and images were analyzed with the Zen 2.6 software.

### Behavior Tests

The tests were realized in isolators in a Biosafety level-3 facility that were specially equipped. We performed 3 sessions of behavioral tests starting at 14, 28 and 76 dpi. At each session, the tests were performed always in the same order, one test/day: light/dark box test (14, 28 and 76 dpi), sucrose splash test (15, 29 and 77 dpi), novelty-suppressed feeding test (16, 30 and 78 dpi), and novel object recognition (17, 31 and 79 dpi). Tests with infected and mock-infected animals were performed concomitantly, in separated BSL-3 isolators.

#### Light/dark box test

This test was based on a published protocol ^92^ with few modifications. The light/dark box apparatus consisted of a box with two compartments with the same size: a dark chamber (black walls with upper lid) and a light chamber (white wall, no upper lid). The chambers, which dimensions were adapted to fit an BSL-3 isolator (32 x 25 x 18 cm) were connected by a door (5 x 5.5 cm) in the middle of the wall. A standard commercial camera (C920 HD Pro, Logitech) was positioned around 30 cm above the white box, connected to a computer. A standard commercial 8W LED lamp was also positioned around 30 cm above the white box to increase lighting. The animals were individually placed in the white box and were allowed to freely explore the light/dark box for 6 min. The boxes were cleaned with 70% ethanol between trials to avoid olfactory stimuli. One main output: the total time spent in the dark chamber, and two secondary outputs: the number of whole-body transitions from one chamber to another and the latency to enter the dark chamber were annotated manually by analyzing the recorded videos. Increased time spent in the dark box corresponds to an anxious behavior.

#### Sucrose splash test

The test was based on an available protocol ^93^. Briefly, the hamsters were isolated in their home cage and sprayed on the dorsal coat with a 10% sucrose solution (3 squirts/animal). The total time grooming was considered the main output of the test, and the latency to start grooming and the number of grooming sessions were considered as secondary outputs. These variables were recorded manually over 5 min using a chronometer. Time grooming corresponds to an index of motivational and self-care behavior and decreased grooming time is considered as a depression-like behavior.

#### Novelty-suppressed feeding test

This test was adapted from a published protocol ^94^. The test was held in a transparent arena (37 x 29 x 18 cm) with clean standard bedding. A filter paper disc (diameter of 12 cm) was placed at the center of the arena, and a food pellet was placed in the middle of the paper disc. A standard commercial 8W LED lamp was positioned around 30 cm above the arena to increase lighting. One day before testing the hamsters were fasted. On the day of the test, the hamsters were individually placed in one corner of the arena. The test lasted a maximum of 10 minutes, and the latency to eat the food (defined as the time to grasp the food pellet with forepaws and bite; and not only approaching or sniffing the food pellet) was recorded using a chronometer. As soon as the food was eaten, the hamsters were removed from the arena and placed back into their home cage with *ad libitum* access to food. If the animal did not eat the food after 10 minutes, the latency to eat was considered to be 10 minutes for statistical purposes. Increased latency to eat indicates hyponeophagia and corresponds to an anxious behavior.

#### Novel object recognition test

This test was adapted from a published protocol ^95^. The test was held in a transparent arena (37 x 29 x 18 cm) with clean standard bedding and consisted of two trials, filmed by a standard commercial (C920 HD Pro, Logitech) positioned around 30 cm above the arena. In trial 1, the animals were allowed to explore the arena with two identical objects placed at an equal distance in the middle of the arena (configuration ●-● or ▲-▲). The objects were 3D-printed green cylinders (diameter 4 cm, height 4 cm) and pink triangular prisms (base 3.8 cm, height 3.3 cm, length 4 cm). To avoid object preference, half of the animals of each experimental group were exposed to two green cylinders and the other half to two pink prisms. Further, the objects were cleaned with 70% ethanol between trials to avoid olfactory stimuli. The animals were placed in a corner of the arena and filmed during 5 min. To assess short-term memory, after 45 min of rest in the home cage, in trial 2, the animals were allowed to explore the same arena, this time containing one familiar object (the same as in trial 1) and one novel object (configuration ●-▲). Consequently, in trial 2, the arenas contained one green cylinder, and one pink prism placed at an equal distance in the middle of the arena. The animals were placed in a corner of the arena and filmed during 5 min. The exploration time (defined as the time sniffing or touching the object, but not sitting on it) was measured manually by analyzing the recorded videos. The time exploring the known object (A) and the novel object (B) was computed and the discrimination index was calculated according to the formula { B–A / B+A }. Decreased discrimination index corresponds to less time spent exploring the new object and is indicative of short-term memory impairment.

### Statistical analysis

Behavioral tests described above were analyzed using mixed-effect model in which a random effect term accounted for the within-individual correlation arising from the longitudinal study design. The fixed effects corresponded to the set of available covariates, *i.e.* hamster sex, SARS-CoV-2 strain and days post-inoculation, along with the corresponding interactions should they significantly modulate the response. These analyses were performed using R v.4.4.1 ^96^ with packages lme4 (v1.1-35.3) ^97^ and emmeans (v1.10.1) ^98^. Other statistical analyses were performed using Prism 10 (GraphPad, version 10.2.3, San Diego).

## Acknowledgements

The SARS-CoV-2 strain was supplied by the National Reference Centre for Respiratory Viruses hosted by Institut Pasteur (Paris, France) and headed by Pr. Sylvie van der Werf. The human sample from which strain 2019-nCoV/IDF0372/2020 was isolated has been provided by Dr. Xavier Lescure and Pr. Yazdan Yazdanpanah from the Bichat Hospital (Paris, France). The isolate SARS-CoV-2 Delta/2021/I7.2 200 (Delta, GISAID ID: EPI_ISL_2029113) and the isolate SARS-CoV-2 Omicron/B.1.1.529 (Omicron/BA.1, GISAID ID: EPI_ISL_6794907) were supplied by the Virus and Immunity Unit hosted by Institut Pasteur and headed by Dr. Olivier Schwartz. This work was supported by the Fondation pour la Recherche Médicale (grant ANRS MIE 202112015304) and by the Institut Pasteur’s Programme Fédérateur de Recherche 4 (PFR-4 – Long Covid). A.C. acknowledges funding from the Institut Pasteur’s 2022-2023 Brain Axis SRA3 M2 Master Student Call. We thank Grégory Inizan for his help in producing customized apparatus for hamster behavioral testing, as well as Gautier Penchinat for 3D-printing the objects used in the novel object recognition test. We would like to thank Marion Berard, Laetitia Breton, Rachid Chennouf, Hamidou Diakhate, Eddie Maranghi and Mathilde Dubot for their help in implementing animal behavior tests in the Institut Pasteur animal facilities. We acknowledge Yakov Vitrenko, Biomics Platform, C2RT, Institut Pasteur, supported by France Génomique (ANR-10-INBS-09) and IBISA. We also thank Esma Karkeni, Single Cell Biomarkers Unit of Technology and Service (scBiomarkers UTechS), Institut Pasteur, for support with the SIMOA assays. Finally, we would like to thank Pr. Sandie Munier for proofreading the manuscript.

## Author Contributions

Conceptualization: A.C., G.D.M., and H.B.

Methodology: G.D.M., F.L., T.O., and E.K.

Investigation: A.C., F.L., L.K., M.T., G.D.M.

Funding Acquisition: G.D.M., and H.B.

Resources: G.D.M., F.L., D.H., and E.K.

Supervision: G.D.M.

Writing – Original Draft: A.C., and G.D.M.

Writing – Review & Editing: all authors

## Data availability

All data associated with this study are present in the paper or the Supplementary Materials. RNA-seq data are available in the following databases: XXX.

## SUPPLEMENTARY FIGURE LEGENDS

**Supplementary Fig. 1. Viral load in the brainstem and serum neurofilament light chain levels of hamsters infected with SARS-CoV-2 Wuhan.** (A) Viral RNA load kinetics in the brainstem of male hamsters. (B) Comparison of serum neurofilament light chain (NfL) levels between mock and Wuhan-infected male hamsters at 80 days post infection (dpi). (C) Viral RNA load kinetics in the brainstem of female hamsters. (B) Comparison of serum neurofilament light chain (NfL) levels between mock and Wuhan-infected female hamsters at 80dpi. (A,C). Viral RNA load kinetics in the brainstem of hamsters infected with Wuhan evaluated at 4 hours, 1, 2, 4, 14, 30 and 80 dpi. Genomic and sub-genomic viral RNA were assessed based on the RdRp and E gene sequence. Gray lines connect symbols from the same animals. mock, 2dpi, 4dpi, 14 dpi and 30 dpi (n=3/group): 4h, 1dpi and 14dpi (n=2/group); 80dpi (n=4/group). Related to Fig.1.

**Supplementary Fig. 2. SARS-CoV-2 distribution in different tissues of hamsters infected with Wuhan or the variants Delta and Omicron/BA.1 at 80 days post infection (dpi).** (A-B) Virus distribution at 80 dpi in different tissues: nasal turbinates, lungs, olfactory bulbs, cerebral cortex and cerebellum infected with SARS-CoV-2 Wuhan, Delta or Omicron/BA.1 in male (A) or female (B) hamsters (n=4/group). Genomic and sub-genomic viral RNA were assessed based on the RdRp and E gene sequence. Gray lines connect symbols from the same animals. Related to Fig.1.

**Supplementary Fig. 3. Histopathological analysis in the brain of hamsters intranasally-infected with SARS-CoV-2 Wuhan.** (A-E) HE staining of the brain of a mock-infected hamster: olfactory bulb, between the olfactory nerve layer and the glomerular layer (A), thalamus (B), midbrain (C), pons (D) and medulla oblongata (subependymal zone; E). (F-J) HE staining of the brain at 4 days post-infection (dpi). (K-O) HE staining of the brain at 80 dpi. No major alteration was observed. Representative images: mock-infected (n=4 males + 4 females), acute COVID-19 (n=4 males), long Covid (n=4 males + 4 females). Scale bars = 100 µm. Related to Fig. 2.

**Supplementary Fig. 4. Histopathological analyses in the airways of hamsters intranasally-infected with SARS-CoV-2 Wuhan.** (A) Formalin-fixed and decalcified macroscopic and submacroscopic views of the nasal cavity of a hamster after a sagittal (upper panels) or a transversal section (bottom panels). The dotted lines in the sagittal view indicate the sections to obtain the transversal view. The dotted square in the submacroscopic transversal view (septum) indicates the region from where the images B-D were obtained. (B-D) Olfactory mucosa from a mock-infected hamster (B), and infected hamsters at 4 days post-infection (dpi; C) and 80 dpi (D). OE: olfactory epithelium, BG: Bowman’s gland, NB: nerve bundles (*filia olfactoria*). Note the extensive destruction of the olfactory epithelium (arrowhead) at 4 dpi. (E-G) Submacroscopic views of the lung of a mock-infected hamster (E), and infected hamsters at 4 dpi (F) and 80 dpi (G). Note an extensive zone of inflammation, congestion and necrosis (arrowhead) at 4 dpi. Representative images: mock-infected (n=4 males + 4 females), acute COVID-19 (n=4 males), long Covid (n=4 males + 4 females). Scale bars: A macroscopic views = 5 mm; sub-macroscopic views = 1 mm, B-D = 100 µm, E-G = 1 mm. Related to Fig. 2.

**Supplementary Fig. 5. Intranasal SARS-CoV-2 infection alters the brainstem innate immune profile in hamsters.** (A-D) Validation targets in the brainstem of male and female hamsters intranasally-infected with Wuhan or the variants Delta and Omicron/BA.1 at 4 and 80 dpi (n=4/group) compared to mock-infected (n=3/group at 4dpi and n=4/group at 80 dpi). Kruskal-Wallis test (the adjusted is *p* value is indicated if *p* < 0.1 and in bold if *p* < 0.05).

**Supplementary Fig. 6. Intranasal SARS-CoV-2 infection alters the brainstem synaptic transcriptome profile in hamsters.** (A-J) Validation targets in the brainstem of male and female hamsters intranasally-infected with Wuhan or the variants Delta and Omicron/BA.1 at 4 and 80 dpi (n=4/group) compared to mock-infected (n=3/group at 4dpi and n=4/group at 80 dpi). Kruskal-Wallis test (the adjusted is *p* value is indicated if *p* < 0.1 and in bold if *p* < 0.05). Related to Fig. 4.

**Supplementary Fig. 7. Intranasal SARS-CoV-2 infection induces a neurodegenerative profile in hamsters.** (A-D) Validation targets in the brainstem of male and female hamsters intranasally-infected with Wuhan or the variants Delta and Omicron/BA.1 at 4 and 80 dpi (n=4/group) compared to mock-infected (n=3/group at 4dpi and n=4/group at 80 dpi). Kruskal-Wallis test (the adjusted is *p* value is indicated if *p* < 0.1 and in bold if *p* < 0.05). Related to Fig. 5.

**Supplementary Fig. 8. Long-term impact of SARS-CoV-2 infection on anxiety-like behavior in hamsters.** (A-B) Follow-up of the novelty-suppressed feeding test assessed at 16-, 30- and 78-days post infection (dpi) in male (A) and female (B) hamsters. Increased latency to eat indicates hyponeophagia and corresponds to an anxious behavior. Horizontal dotted lines indicate the upper 95% confidence limit of the median. Kruskal-Wallis test (the adjusted is *p* value is indicated if *p* < 0.1 and in bold if *p* < 0.05). Related to Fig. 6.

**Supplementary Fig. 9. Long-term impact of SARS-CoV-2 infection on anxiety-like behavior in hamsters.** (A) Follow-up of the light-dark box test assessed at 14-, 28- and 76-days post infection (dpi) in male hamsters. Increased time spent in the dark box corresponds to an anxious behavior. (B) Average predicted time in the dark analyzed by mixed-models regression in male hamsters. (C) Follow-up of the light-dark box test assessed at 14, 28 and 76 dpi in female hamsters. (D) Average predicted time in the dark analyzed by mixed-models regression in female hamsters. Horizontal dotted lines indicate the upper 95% confidence limit of the median. Kruskal-Wallis test (the adjusted is *p* value is indicated if *p* < 0.1 and in bold if *p* < 0.05). In the mixed-models regression, the line indicates the mean with its 95% confidence interval. (E-H) Secondary outputs of the test. Latency to enter in the dark chamber (E,F) and total number of transitions between the light and the dark chambers (G,H). Related to Fig. 6.

**Supplementary Fig. 10. Long-term impact of SARS-CoV-2 infection on depression-like behavior in hamsters.** (A-B) Follow-up of the sucrose splash test assessed at 15-, 29- and 77-days post infection (dpi) in male (A) and female (B) hamsters. Time grooming corresponds to an index of motivational and self-care behavior and decreased grooming time is considered as a depression-like behavior. (C-D) Latency to start grooming at 15, 29 and 77 days post-infection (dpi) in male (C) and female hamsters (D). (E-F) Total number of grooming sessions at 15, 29 and 77 dpi in male hamsters (E) and female (F) hamsters. Horizontal dotted lines indicate the upper (C,D) or the lower (A,B,E,F) 95% confidence limit of the median. Kruskal-Wallis test (the adjusted is *p* value is indicated if *p* < 0.1 and in bold if *p* < 0.05). Related to Fig. 6.

**Supplementary Fig. 11. Long-term impact of SARS-CoV-2 infection on recognition memory in hamsters.** (A-B) Follow-up of the novel object recognition test assessed at 17-, 31- and 79-days post infection (dpi) in male (A) and female (B) hamsters. Decreased discrimination index corresponds to less time spent exploring the new object and is indicative of short-term memory impairment. Horizontal dotted lines indicate the lower 95% confidence limit of the median. (C-D) Follow-up of exploration time for identical objects (●-● or ▲-▲) in the novel object recognition test assessed at 17-, 31- and 79- dpi in male (C) and female (D) hamsters. (E-F) Follow-up of exploration time for old and novel objects (●-▲) in the novel object recognition test assessed at 17-, 31- and 79- dpi in male (E) and female (F) hamsters. Kruskal-Wallis test (the adjusted is *p* value is indicated if *p* < 0.1 and in bold if *p* < 0.05). Related to Fig. 6,

**Supplementary Table 1.**
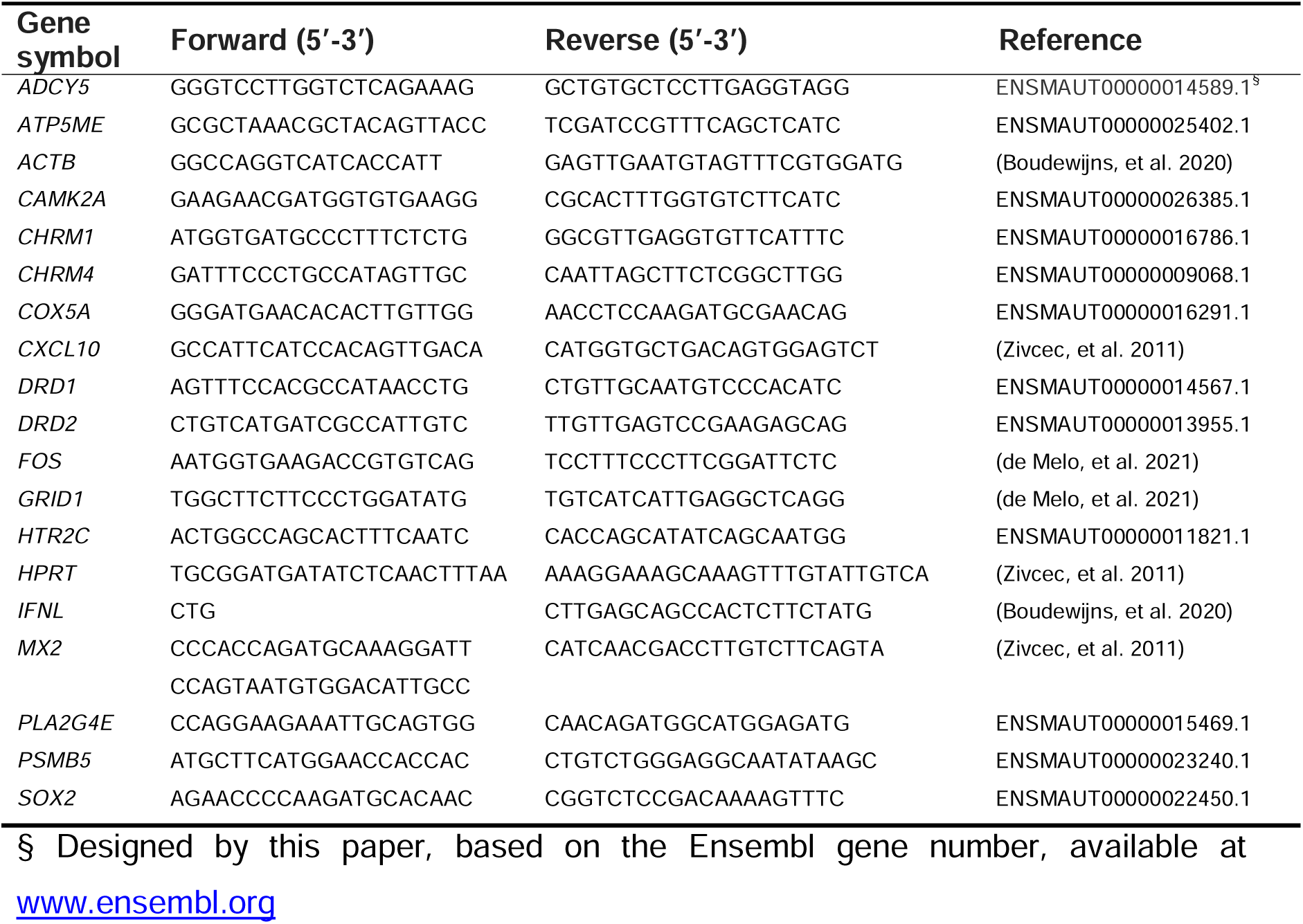
Primer sequences used for qPCR in the golden hamster tissues. **References of Supplementary Table 1** Boudewijns, *et al.* STAT2 signaling restricts viral dissemination but drives severe pneumonia in SARS-CoV-2 infected hamsters. *Nat Commun* 11, 5838 (2020). de Melo, G. D. *et al.* Attenuation of clinical and immunological outcomes during SARS-CoV-2 infection by ivermectin. *EMBO Mol Med* 13, e14122 (2021). Zivcec, *et al*. Validation of assays to monitor immune responses in the Syrian golden hamster (Mesocricetus auratus*). J Immunol Methods* (2011).

